# Alpha-defensin binding expands human adenovirus tropism

**DOI:** 10.1101/2024.05.30.596681

**Authors:** Cheng Zhao, Jessica M. Porter, Phillip C. Burke, Niklas Arnberg, Jason G. Smith

**Author notes:** Address correspondence to Jason G. Smith.

## Abstract

Mammalian α-defensins are a family of abundant effector peptides of the mucosal innate immune system. Although primarily considered to be antimicrobial, α-defensins can increase rather than block infection by certain prominent bacterial and viral pathogens in cell culture and *in vivo*. We have shown previously that exposure of mouse and human adenoviruses (HAdVs) to α-defensins is able to overcome competitive inhibitors that block cell binding, leading us to hypothesize a defensin-mediated binding mechanism that is independent of known viral receptors. To test this hypothesis, we used genetic approaches to demonstrate that none of several primary receptors nor integrin co-receptors are needed for human α-defensin-mediated binding of HAdV to cells; however, infection remains integrin dependent. Thus, our studies have revealed a novel pathway for HAdV binding to cells that bypasses viral primary receptors. We speculate that this pathway functions in parallel with receptor-mediated entry and contributes to α-defensin-enhanced infection of susceptible cells. Remarkably, we also found that in the presence of α-defensins, HAdV tropism is expanded to non-susceptible cells, even when viruses are exposed to a mixture of both susceptible and non-susceptible cells. Therefore, we propose that in the presence of sufficient concentrations of α-defensins, such as in the lung or gut, integrin expression rather than primary receptor expression will dictate HAdV tropism *in vivo*. In summary, α-defensins may contribute to tissue tropism not only through the neutralization of susceptible viruses but also by allowing certain defensin-resistant viruses to bind to cells independently of previously described mechanisms.

**Author Summary:** In this study, we demonstrate a novel mechanism for binding of human adenoviruses (HAdVs) to cells that is dependent upon interactions with α-defensin host defense peptides but is independent of known viral receptors and co-receptors. To block normal receptor-mediated HAdV infection, we made genetic changes to both host cells and HAdVs. Under these conditions, α-defensins restored cell binding; however, infection still required the function of HAdV integrin co-receptors. This was true for multiple types of HAdVs that use different primary receptors and for cells that are either naturally devoid of HAdV receptors or were engineered to be receptor deficient. These observations suggest that in the presence of concentrations of α-defensins that would be found naturally in the lung or intestine, there are two parallel pathways for HAdV binding to cells that converge on integrins for productive infection. Moreover, these binding pathways function independently, and both operate in mixed culture. Thus, we have found that viruses can co-opt host defense molecules to expand their tropism.

## Introduction

Mammalian defensins are host defense peptide effectors of the innate immune system [1, 2]. Humans abundantly express two subtypes, α– and β-defensins. α-defensins can be further categorized as myeloid or enteric based on their expression patterns and gene organization. The human myeloid α-defensins, human neutrophil peptide 1 to 4 (HNP1-4), are produced primarily by neutrophils. The human enteric α-defensins, human defensin 5 and 6 (HD5 and HD6), are constitutively expressed and secreted by specialized epithelial cells of the small intestine and in the genitourinary tract. Both α– and β-defensins can inactivate Gram-positive and –negative bacteria as well as some types of enveloped viruses through a variety of mechanisms [3]. α-defensins have the additional capacity to neutralize non-enveloped viruses from several families including *Adenoviridae, Papillomaviridae, Polyomaviridae, Parvoviridae*, and *Reoviridae* [4–7]. Thus, mammalian defensins are broadly antimicrobial.

Although the antimicrobial activities of defensins have been investigated most extensively, particular types of human adenovirus (HAdV), adeno-associated virus (AAV), mouse adenovirus (MAdV), rotavirus, and human immunodeficiency virus have been shown to be either resistant to α-defensin neutralization or even able to commandeer α-defensins to enhance their infection [6–13]. Similarly, infection by the bacterial pathogen *Shigella* is increased by α-defensins, and some commensal bacterial species are resistant to α-defensin killing [14–18]. These observations suggest an evolutionary adaptation of certain microbes upon exposure to α-defensins during replication or transmission, and α-defensins have been shown to exert sufficient selective pressure for the evolution of viruses in cell culture [8]. However, mechanisms of viral resistance to or appropriation of α-defensins are poorly understood.

In this study, we used HAdVs to investigate how viruses hijack α-defensins. HAdV infection is initiated by interactions between viral proteins that comprise the non-enveloped, icosahedral capsid and host receptors. All three major capsid proteins, fiber, penton base, and hexon, have been shown to play a role in cell attachment [19, 20]. In cell culture, three primary receptors that bind with high affinity to the distal knob domain of fiber have been identified: coxsackievirus and adenovirus receptor (CAR), CD46, and desmoglein-2 [19]. In addition, lower affinity interactions between the fiber knob and polysialic acids linked to glycoproteins and gangliosides mediate attachment of some HAdVs. HAdVs are classified into species A-G and then further categorized into over 110 serotypes and genotypes [21, 22]. In most cases a particular HAdV serotype binds to only one of these receptors; however, there are some exceptions [23]. Moreover, other modes of attachment, such as a direct interaction between the hexon of certain HAdV-D serotypes and CD46, have been uncovered [20]. Upon cell attachment, most HAdVs are internalized upon signaling initiated by interactions between a conserved arginine-glycine-aspartate (RGD) motif in penton base and any of several RGD-binding integrin co-receptors [19, 24]. *In vivo*, additional attachment mechanisms have been described in which host proteins such as coagulation proteins (e.g., factor X) or the iron regulatory protein lactoferrin bind to hexon and bridge capsid interactions with the cell [25–27]. Therefore, although HAdV entry has been extensively studied in cell culture, how the complexity of interactions dictates tissue tropism *in vivo* is less well understood.

We provide evidence that α-defensin expression may contribute to HAdV tissue tropism. In prior studies we found that α-defensins allowed certain serotypes of both HAdVs and MAdVs to overcome competitive inhibition of receptor binding to successfully infect cells [9, 10]. Moreover, α-defensin interactions with all HAdVs led to increased cell binding [8, 10, 28]. Together, these data suggest that α-defensins may influence how AdVs bind to cells. In this study, we used biochemical, genetic, and enzymatic approaches to demonstrate that HAdVs in complex with α-defensins can attach to cells independently of known viral receptors, although infection remains integrin-dependent. We postulate that this alternative binding mechanism at least in part explains α-defensin-enhanced infection of some HAdV serotypes; however, we also show that this interaction expands the tropism of HAdV, implicating integrin expression rather than primary receptor expression as a major determinant of tropism. Thus, α-defensins may contribute to tissue tropism not only through the neutralization of susceptible serotypes during entry but also by allowing certain serotypes to bind to cells independently of previously described mechanisms.

## Results

### HD5 promotes sialic acid-independent infection of human epithelial cells by HAdV-D64

Our prior studies of HAdV demonstrated that α-defensin binding to the viral capsid allows infection to proceed despite competition for primary receptor binding by a molar excess of fiber knob, suggesting that α-defensins mediate a viral receptor-independent route of entry [10]. An alternative explanation for this observation is that α-defensin interactions allow recombinant fiber knob to bridge an interaction between the virus and the cell. A similar mechanism has been described for HAdV-D37 fiber knob-mediated infection of myeloid cells by HAdV-C5 [29]. To determine whether the ability of α-defensins to overcome a competitive inhibitor is unique to fiber knob, we sought to mask a HAdV primary receptor with a non-viral competitor. We chose HAdV-D64 for these experiments for two reasons. First, like all HAdV-D serotypes that we have tested, HAdV-D64 is naturally resistant to neutralization by human α-defensins, as evidenced by an increase in eGFP expression from A549 cells transduced by a HAdV-D64-based vector in the presence of 2.5 µM or 5 µM HD5 compared to control (Fig 1A) [8, 10]. This is unlike HAdV-C5, which is neutralized by HD5 (Fig 1A) [8, 10, 30, 31]. Second, HAdV-D64 is closely related to other epidemic keratoconjunctivitis (EKC)-causing HAdV-D serotypes that use sialic acid as their primary receptors [19, 32]. Binding to sialic acid can be potently inhibited by the lectin wheat germ agglutinin (WGA), and consistent with the use of sialic acid as a receptor, HAdV-D64 infection of A549 cells in the absence of the human enteric α-defensin HD5 was reduced by WGA treatment to ∼20% (Fig 1B). However, in the presence of increasing concentrations of HD5, infection of WGA treated cells was restored in a dose-dependent fashion to levels equal to or exceeding those of untreated cells. Thus, the ability of α-defensins to overcome inhibition of the primary receptor interaction is not restricted to competition with recombinant viral proteins.

**Figure 1.**
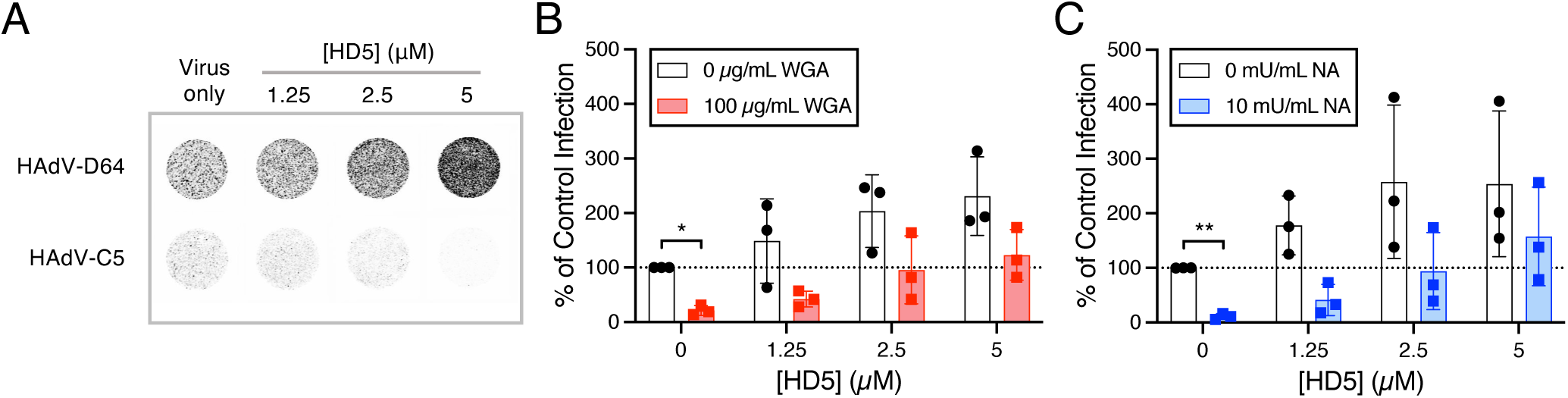
HD5 promotes HAdV-D64 infection despite blockade of the primary receptor, sialic acid. (A) Representative images of A549 cell monolayers in 96-well plates infected with HAdV-C5 or HAdV-D64 in the presence or absence of HD5 were obtained at a resolution of 50 μm. Grayscale intensity correlates with eGFP expression. (B–C) HAdV-D64 infection of A549 cells that were treated or not with (B) 100 µg/mL WGA or (C) 10 mU/mL neuraminidase (NA) in the presence or absence of HD5. Each point is an independent experiment, and bars are the mean ± SD of the percent infectivity compared to control, untreated cells infected in the absence of HD5 (100%).

Rather than mediating receptor-independent cell binding, α-defensins could stabilize the fiber-receptor interaction, thereby increasing the avidity of the interaction. To test this hypothesis, instead of adding excess inhibitor, we sought to remove the receptor from the cell surface by treating cells prior to infection with neuraminidase, an enzyme that cleaves sialic acids from proteins and lipids. As was true for WGA, neuraminidase treatment reduced HAdV-D64 infection in the absence of HD5 to ∼10% of the level of infection of untreated cells (Fig 1C). HD5 again restored infection to levels greater than or equal to untreated cells. Taken together, these results suggest that HD5 promotes cell binding and infection by HAdV in the absence of a primary receptor. They also provide evidence against a specific role for α-defensin-stabilized interactions either between recombinant fiber knob and the capsid or between fiber and the primary receptor.

### HD5-mediated binding of HAdV to cells is independent of known viral receptors

Although exposure to HD5 was able to overcome masking or enzymatic removal of the primary HAdV-D64 receptor, the effect of these treatments was incomplete, as 10–20% of infection remained after treatment (Fig 1B and C). Moreover, other receptors (e.g., CD46) have been postulated for EKC-causing HAdVs [33]. Thus, we cannot formally exclude a role for primary viral receptors in α-defensin-mediated infection from our previous experiments. To address these shortcomings, we turned to an eGFP-expressing HAdV-5-based vector (C5/D64-HVR1) that uses coxsackie-adenovirus receptor (CAR) for initial attachment and has been engineered to be HD5-resistant by replacing the hexon hypervariable region 1 (HVR1) of HAdV-C5 with that of HAdV-D64 (Fig 2A). We then used CRISPR/Cas9 to generate CAR-deleted A549 cells (CAR KO). Successful ablation of CAR was validated by sequencing (Fig 2B) and flow cytometry (Fig 2C), and the specificity of this genetic change for CAR-utilizing viruses was confirmed by infection (Fig 2D). This combination of CAR KO cells and the C5/D64-HVR1 virus allowed us to analyze HD5-mediated infection in the complete absence of known primary viral receptors.

**Figure 2.**
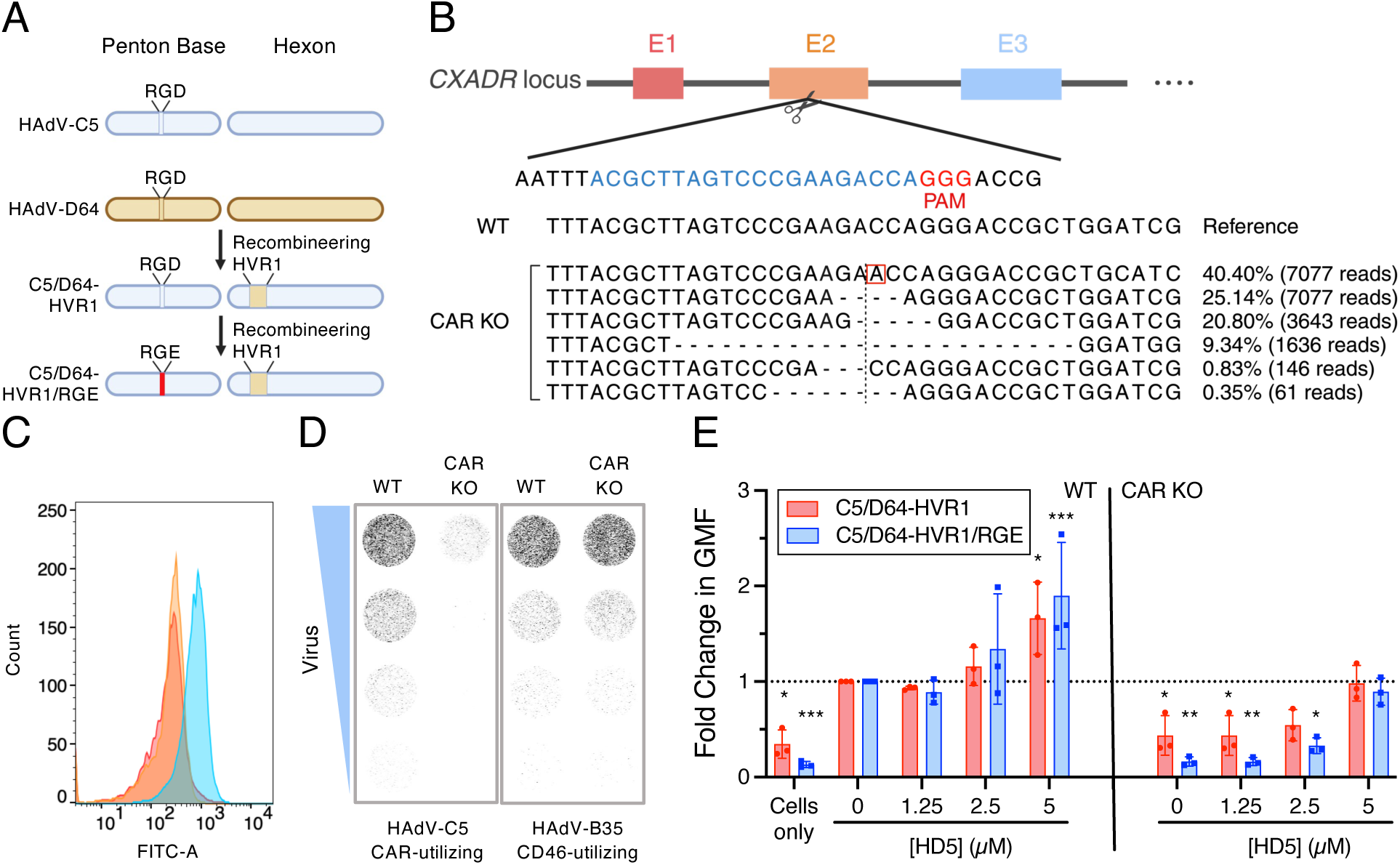
HD5-mediated binding of HAdV to cells is independent of known viral receptors. (A) Construction of C5/D64-HVR1/RGE. Capsid proteins are depicted for HAdV-C5 (blue) and HAdV-D64 (brown). The variable residues in penton base (RGD and RGE) and hexon hypervariable region 1 (HVR1) are indicated. (B) Strategy for editing *CXADR* exon 2 in A549 cells. The small-guide RNA (sgRNA) target site (blue) and proto-spacer adjacent motif (PAM, red) sequences are highlighted. Below are the WT reference allele and an analysis of alleles from the polyclonal CAR KO A549 cells used in this study. The percentage and number of reads for each allele are indicated on the right. The vertical dotted line is the predicted cleavage site, dashes indicate deletions, and the red box denotes an insertion. (C) Validation of CAR KO by flow cytometry analysis. The yellow line represents cells stained with the isotype control, the blue line represents WT cells stained with the anti-CAR antibody, and the red line represents CAR KO cells stained with the anti-CAR antibody. The figure is representative of three independent experiments with similar results. (D) Representative cell monolayers in 96 well plates of WT A549 and CAR KO A549 cells infected with three-fold serial dilutions of HAdV-C5 and HAdV-B35. Images were obtained at a resolution of 50 μm, and grayscale intensity correlates with eGFP expression. (E) Binding of AF488-labeled viruses to A549 or CAR KO A549 cells in the presence of HD5. Data are the fold change in geometric mean fluorescence (GMF) of the entire cell population relative to the no HD5 condition for each virus bound to WT cells. Each point is an independent experiment, and bars are the mean ± SD.

A second caveat to the HAdV-D64 experiments is that following primary attachment, most HAdVs bind to integrin co-receptors via a conserved RGD motif in the penton base protein [19]. This interaction can also serve for initial cell attachment at high multiplicity of infection in the absence of fiber (and CAR, Fig 2D) [34]. Because multiple RGD-binding integrins are expressed in most cells, a knockout strategy is impractical [35]. Therefore, to disrupt the co-receptor interaction, we mutated the RGD motif in the penton base of C5/D64-HVR1 to RGE (Fig 2A), a change that is known to prevent integrin binding [36], to create C5/D64-HVR1/RGE.

With these tools in hand, we were poised to determine whether HD5 could mediate HAdV attachment to cells in the absence of all known receptor interactions. We fluorescently labeled the C5/D64-HVR1 and C5/D64-HVR1/RGE viruses with Alexa Fluor 488 and quantified total cell fluorescence by flow cytometry as a measure of virus binding in the presence and absence of physiologic concentrations of purified HD5. In WT A549 cells, HD5 treatment led to increased binding of both viruses (Fig 2E), which was dose responsive. Consistent with a major role for the fiber/CAR interaction in cell binding, both viruses failed to bind to CAR KO A549 cells in the absence of HD5. However, binding to CAR KO cells was restored for both viruses in the presence of 5 µM HD5 to levels equivalent to their binding to WT cells. Together, these data formally demonstrate that α-defensin-mediated binding of HAdV to cells is independent of known viral receptors.

### **α**-defensin-mediated infection remains dependent on integrin co-receptors

Cell binding is not sufficient for HAdV entry, and multiple signaling pathways that facilitate clathrin-mediated endocytosis and subsequent uncoating in the endosome have been described [37]. To determine if α-defensin-mediated binding leads to productive infection, we treated the C5/D64-HVR1 and C5/D64-HVR1/RGE viruses with or without increasing concentrations of HD5, added them to either WT or CAR KO A549 cells, and quantified infection 24 h p.i. by eGFP expression. An eGFP-expressing vector based on WT HAdV-C5 was used as a control. In the absence of HD5, all three viruses were able to infect WT but not CAR KO cells (Fig 3). In the presence of HD5 on WT cells, WT HAdV-C5 was inhibited in a dose-dependent fashion with an IC_50_ < 2.5 µM. In contrast, infection by C5/D64-HVR1 was moderately increased at HD5 concentrations > 2.5 µM. Infection by C5/D64-HVR1/RGE was resistant up to 10 µM HD5 then began to trend towards neutralization, with a 35% reduction at 20 µM HD5. However, on CAR KO cells, we observed robust infection only for C5/D64-HVR1, which in the presence of > 10 µM HD5 exceeded the level of infection of WT cells by this virus in the absence of HD5. The effect of HD5 on WT HAdV-C5 was biphasic, and infection was only observed at the incompletely neutralizing concentration of 2.5 µM. Remarkably, infection of CAR KO cells by C5/D64-HVR1/RGE was only rescued by HD5 to a low level, ∼1.4% infection at ≥ 10 µM HD5. These data suggest that HD5-mediated HAdV infection still requires integrin co-receptors to trigger internalization following attachment.

**Figure 3.**
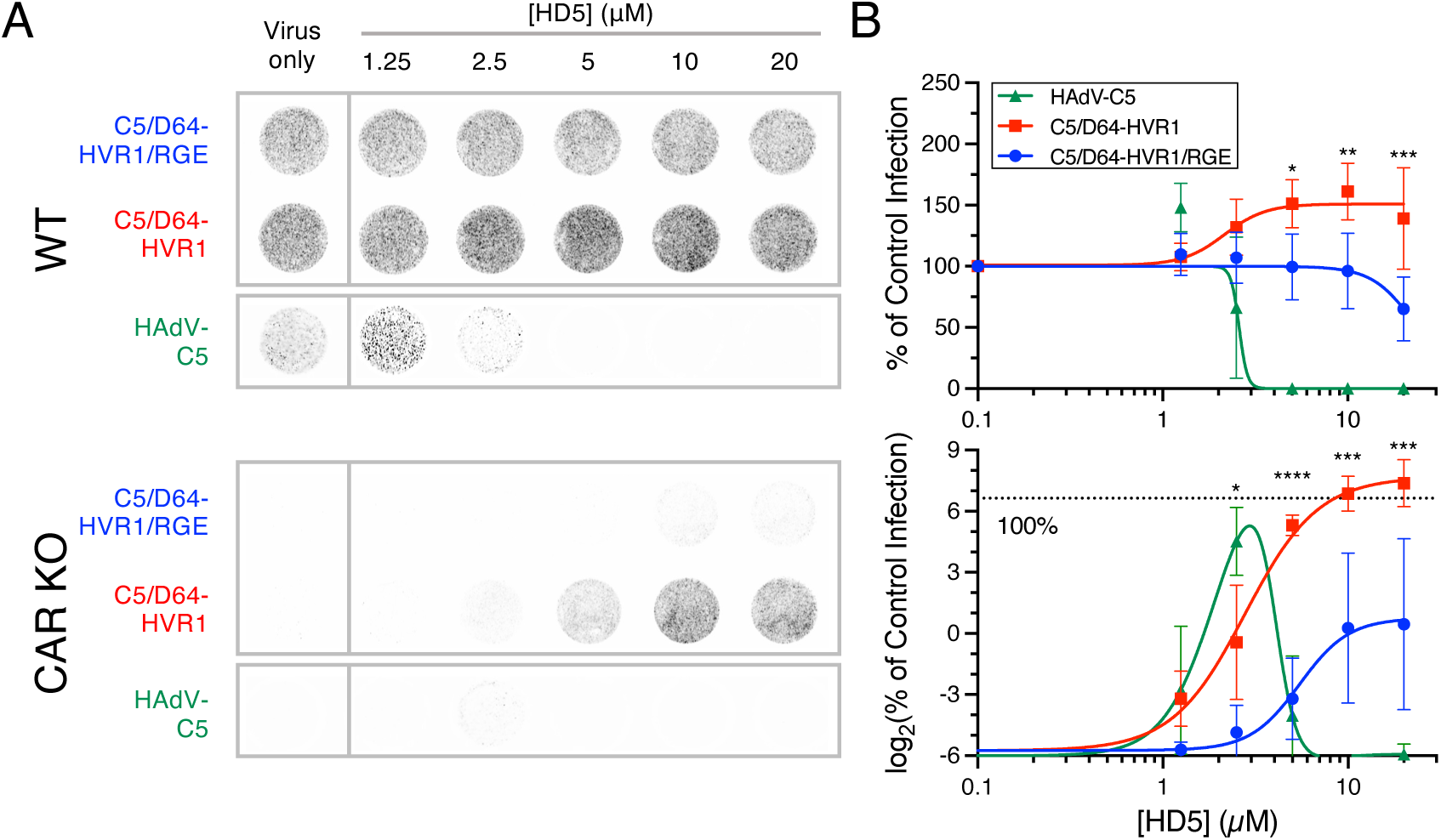
HD5-mediated infection requires integrin co-receptors. (A) Representative images and (B) quantification of infection of (top) A549 and (bottom) CAR KO cells with the indicated viruses in the presence or absence of HD5. Images in A were obtained at a resolution of 50 μm, and grayscale intensity correlates with eGFP expression. In B, data in both graphs are normalized to infection of A549 cells in the absence of defensin (control infection) and are the means ± SD of at least 3 individual experiments.

We next probed the requirement for HD5 self-association in α-defensin-mediated infection. α-Defensins have been shown to multimerize into dimers or oligomers in solution under physiological conditions [38–40], and this multimerization is critical for antiviral activity [6, 28]. Replacement of glutamate 21 with *N*-methyl-glutamate disrupts hydrogen bonding at the dimer interface, resulting in an obligate monomer form of HD5 (E21me HD5) [40]. To investigate the role of HD5 multimerization in receptor-independent infection, we measured the effect of E21me HD5 on both the binding and infectivity of C5/D64-HVR1 and C5/D64-HVR1/RGE. Unlike WT HD5 (Fig 2E), E21me HD5 was unable to rescue cell binding (Fig 4A) or completely restore infectivity (Fig 4B and D) of either virus on CAR KO cells. Rather, only a low level of infection (< 4%) was observed for C5/D64-HVR1 with 20 µM E21me HD5, and no infection (< 0.1%) was observed for C5/D64-HVR1/RGE. Moreover, only C5/D64-HVR1 infection was moderately enhanced on WT cells (Fig 4C). In summary, as was true for neutralization of sensitive viruses, multimerization is critical for efficient α-defensin-mediated infection by defensin-resistant HAdVs.

**Figure 4.**
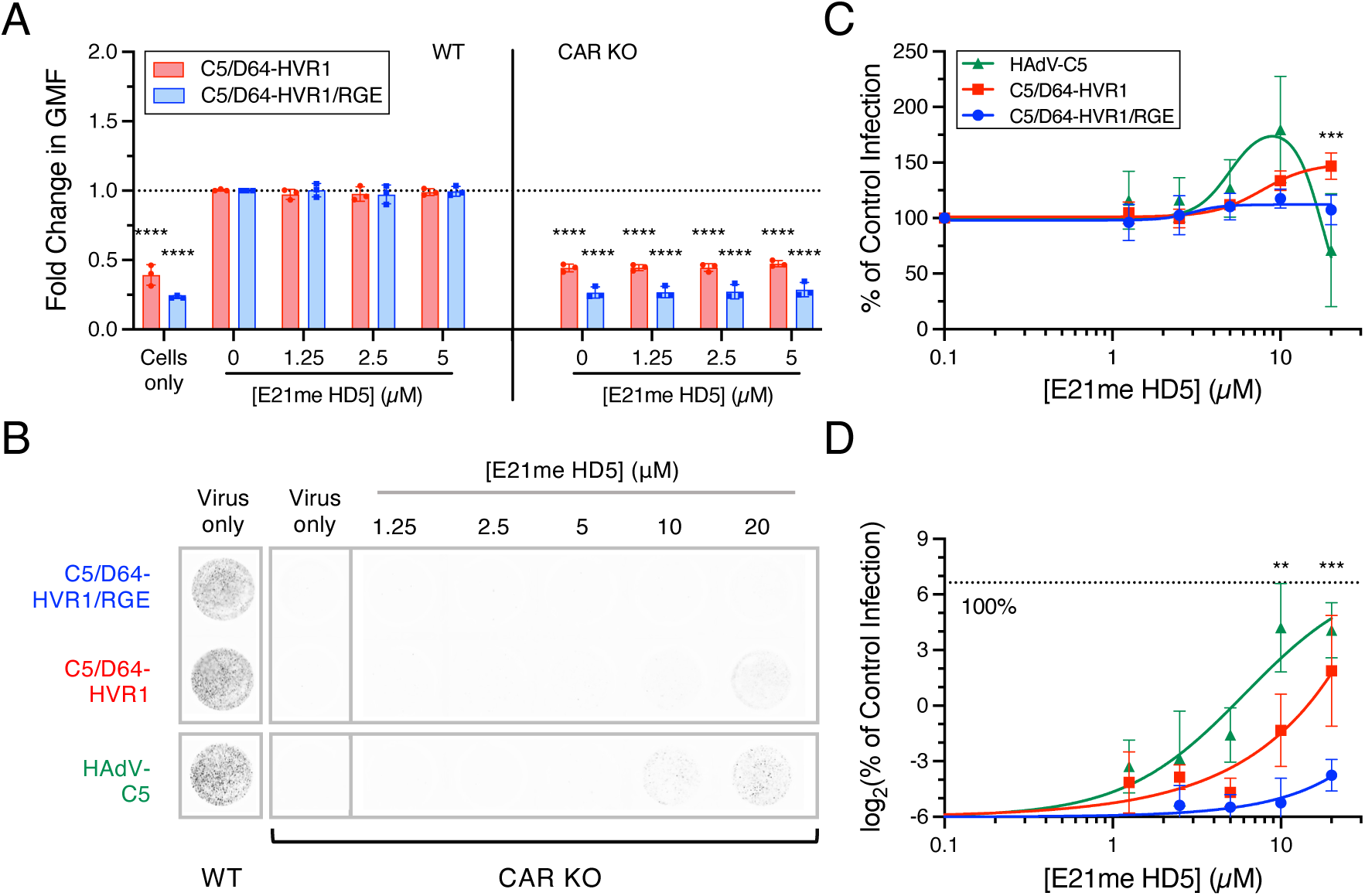
HD5-mediated infection requires HD5 multimerization. (A) Binding of AF488-labeled viruses to A549 or CAR KO cells in the presence of E21me HD5. Data are the fold change in geometric mean fluorescence (GMF) of the entire cell population relative to the no HD5 condition for each virus bound to WT cells. Each point is an independent experiment, and bars are the mean ± SD. (B) Representative images of A549 and CAR KO cells infected with the indicated viruses in the presence or absence of E21me HD5. Images were obtained at a resolution of 50 μm, and grayscale intensity correlates with eGFP expression. Quantification of infection of (C) A549 and (D) CAR KO cells with the indicated viruses in the presence or absence of E21me HD5. Data in both graphs are normalized to infection of A549 cells in the absence of defensin (control infection) and are the means ± SD of at least 3 individual experiments.

To determine if defensin-mediated infection extends to another α-defensin, we assayed HNP1, a myeloid α-defensin that is also capable of enhancing HAdV-D serotypes [10]. On WT cells, HNP1 is not as potent as HD5 in inhibiting WT HAdV-C5 infection when incubated with virus prior to cell attachment (Fig 5), as we described previously [41]. Infection by C5/D64-HVR1 was moderately increased at HNP1 concentrations ≥ 10 µM, while infection by C5/D64-HVR1/RGE was resistant at all HNP1 concentrations. On CAR KO cells, HNP1 resembled HD5 and was able to rescue infection by C5/D64-HVR1 but not C5/D64-HVR1/RGE at concentrations > 5 µM. HNP1 also had a modest ability to rescue WT HAdV-C5 at concentrations of 5–10 µM. Thus, defensin-mediated infection of cells in the absence of primary receptors is a property of multiple human α-defensins and requires both defensin multimerization and the function of integrin co-receptors.

**Figure 5.**
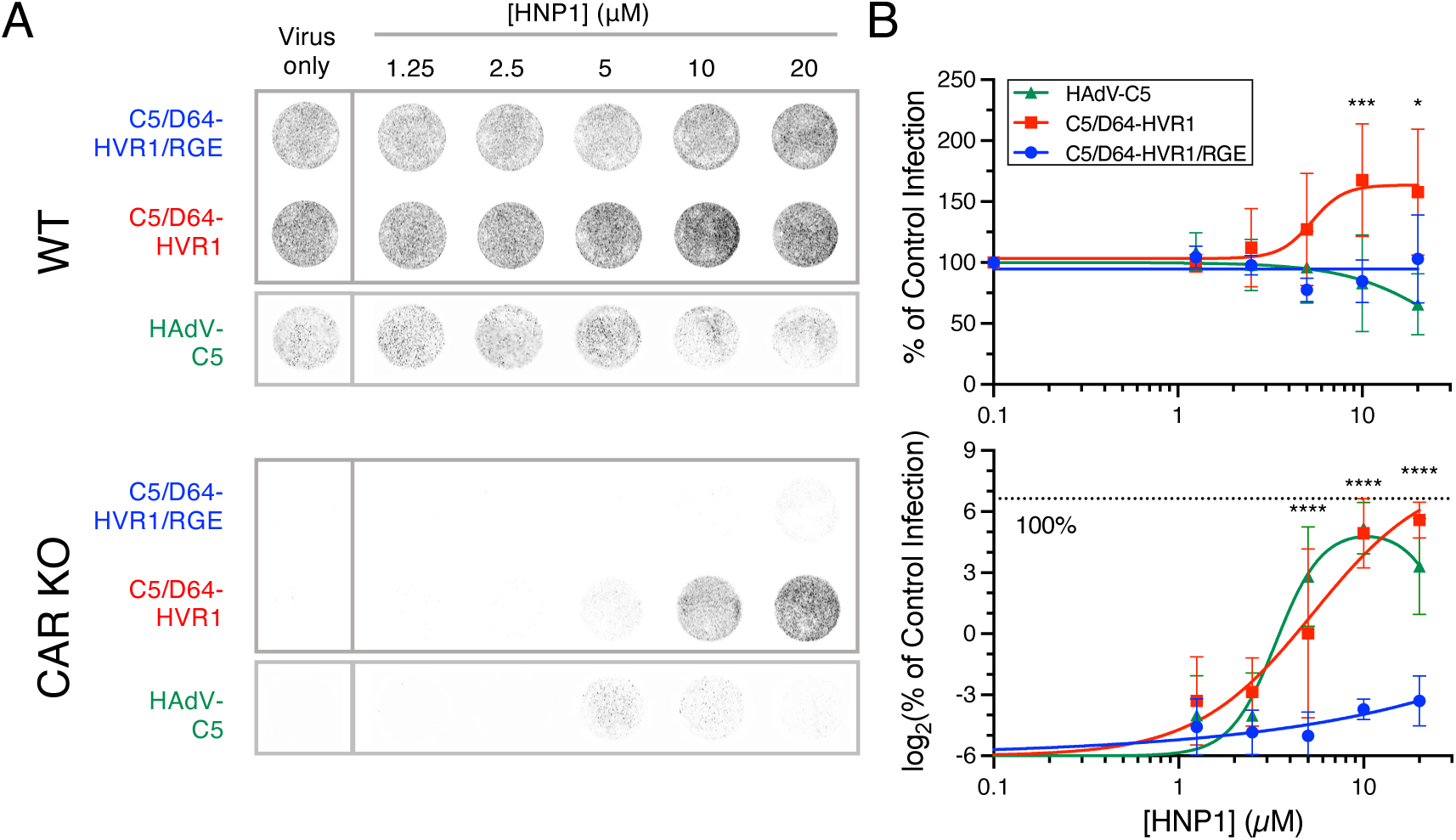
α-Defensin-mediated infection is not restricted to HD5. (A) Representative images and (B) quantification of infection of (top) A549 and (bottom) CAR KO cells with the indicated viruses in the presence or absence of HNP1. Images in A were obtained at a resolution of 50 μm, and grayscale intensity correlates with eGFP expression. In B, data in both graphs are normalized to infection of A549 cells in the absence of defensin (control infection) and are the means ± SD of at least 3 individual experiments.

### Defensin-dependent binding expands the cellular tropism of HAdVs

We speculated that in a more complex biological environment, secreted α-defensins might allow entry of HAdV into cells that naturally lack known primary receptors, thereby expanding tropism. This could be true in the small intestine, where HD5 has been estimated at 14-70 µM in the human ileal lumen at steady state, with higher concentrations in the lumens of crypts near the site of secretion from Paneth cells [42, 43]. It could also occur in the lung, where HNP1 has been measured at concentrations as high as 2.5 mM under some conditions [44]. To test this hypothesis, we turned to CD46-utilizing HAdVs, since mouse CD46 has low homology to human CD46 and cannot serve as a receptor for HAdVs [45]. Because all naturally occurring HAdV-B serotypes that have been tested that use CD46 are neutralized by HD5 and HNP1 [10], we created an α-defensin-resistant HAdV-B-based vector expressing eGFP following the principles of Diaz et al. [8]. We substituted sequences in three capsid proteins of HAdV-B35 (fiber, hexon, and penton base) with the corresponding amino acids of HAdV-D37 (Fig 6A), which like HAdV-D64 is an EKC-causing virus that is naturally resistant to HD5 and HNP1 [10]. We then ascertained the phenotype of this virus (B35/D37-HVR1/Vertex) compared to the parental viruses on A549 cells after pre-incubation with HD5 (Fig 6B). As expected, HAdV-B35 infection was neutralized with an IC_50_ = 4.9 µM, while WT HAdV-D37 was resistant. Consistent with our previous studies identifying capsid protein determinants of α-defensin interactions [8, 10], B35/D37-HVR1/Vertex was enhanced by ≥ 10 µM HD5. Moreover, as was true for C5/D64-HVR1, HD5 rescued the infectivity of this virus on haploid human cells (HAP1) engineered to be devoid of the primary receptor (Fig 6C and D). Thus, we have created and validated a virus suitable for testing our hypothesis.

**Figure 6.**
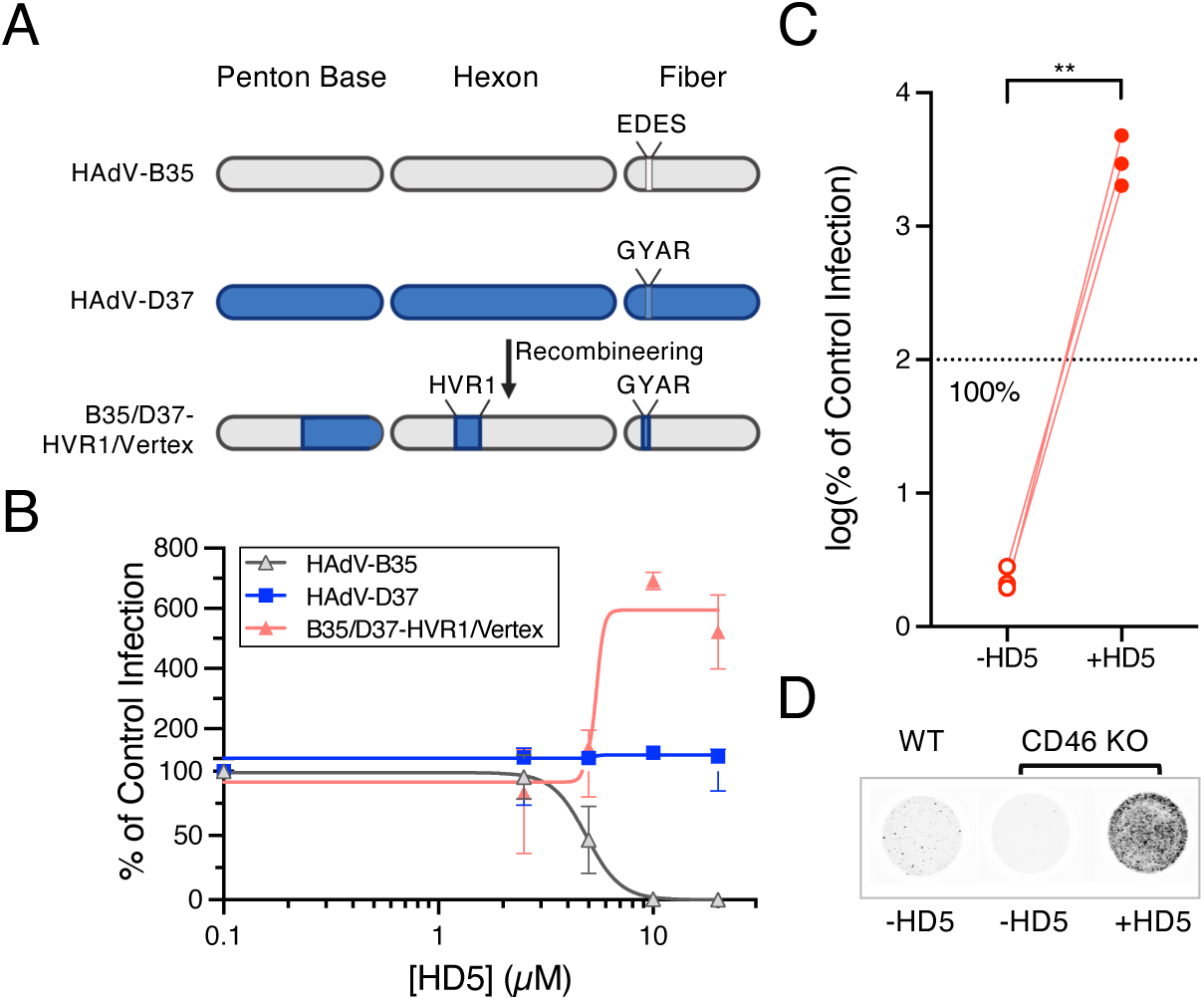
Generation and validation of an HD5-resistant, HAdV-B-based virus. (A) Construction of B35/D37-HVR1/Vertex. Capsid proteins are depicted for HAdV-B35 (gray) and HAdV-D37 (blue). Hexon hypervariable region 1 (HVR1) and the variable residues in fiber (EDES and GYAR) are indicated. (B) Infection of A549 cells with the indicated viruses in the presence or absence of HD5. Data are normalized to infection in the absence of defensin (control infection) and are the means ± SD of at least 3 individual experiments. (C) Infection of CD46 KO HAP1 cells in the presence or absence of 20 µM HD5. Data is normalized to infection of control HAP1 cells in the absence of defensin. Each connected pair of points is an independent experiment. The dashed line is 100% (control) infection. (D) Representative images of the data in C were obtained at a resolution of 50 μm, and grayscale intensity correlates with eGFP expression.

To create a system with multiple cell types, we chose human A549 cells, which are naturally susceptible and permissive to CD46-utilizing HAdVs, and mouse CMT-93 cells, which are neither susceptible nor permissive. Because B35/D37-HVR1/Vertex encodes eGFP, we generated mCherry-expressing CMT-93 cells to distinguish them from A549 cells in mixed culture. As expected, infection of A549 cells in monoculture is robust and unaffected by 20 µM HD5 (Fig 7A). In contrast, CMT-93 cells in monoculture are not infectable by B35/D37-HVR1/Vertex in the absence of HD5; however, 17% of cells are infected in the presence of HD5 (Fig 7B). To determine whether the defensin-mediated, receptor-independent entry pathway functions despite the presence of susceptible cells, we infected mixed cultures of A549 and CMT-93 cells with B35/D37-HVR1/Vertex that was either pre-incubated or not with 20 µM HD5. Under these conditions, A549 cells were robustly infected irrespective of HD5 (Fig 7C and E), and CMT-93 cell infection was again enabled by HD5 (Fig 7D and E). Thus, even in the presence of susceptible cells, B35/D37-HVR1/Vertex can still infect cells lacking primary receptors, demonstrating the ability of α-defensins to expand viral tropism.

**Figure 7.**
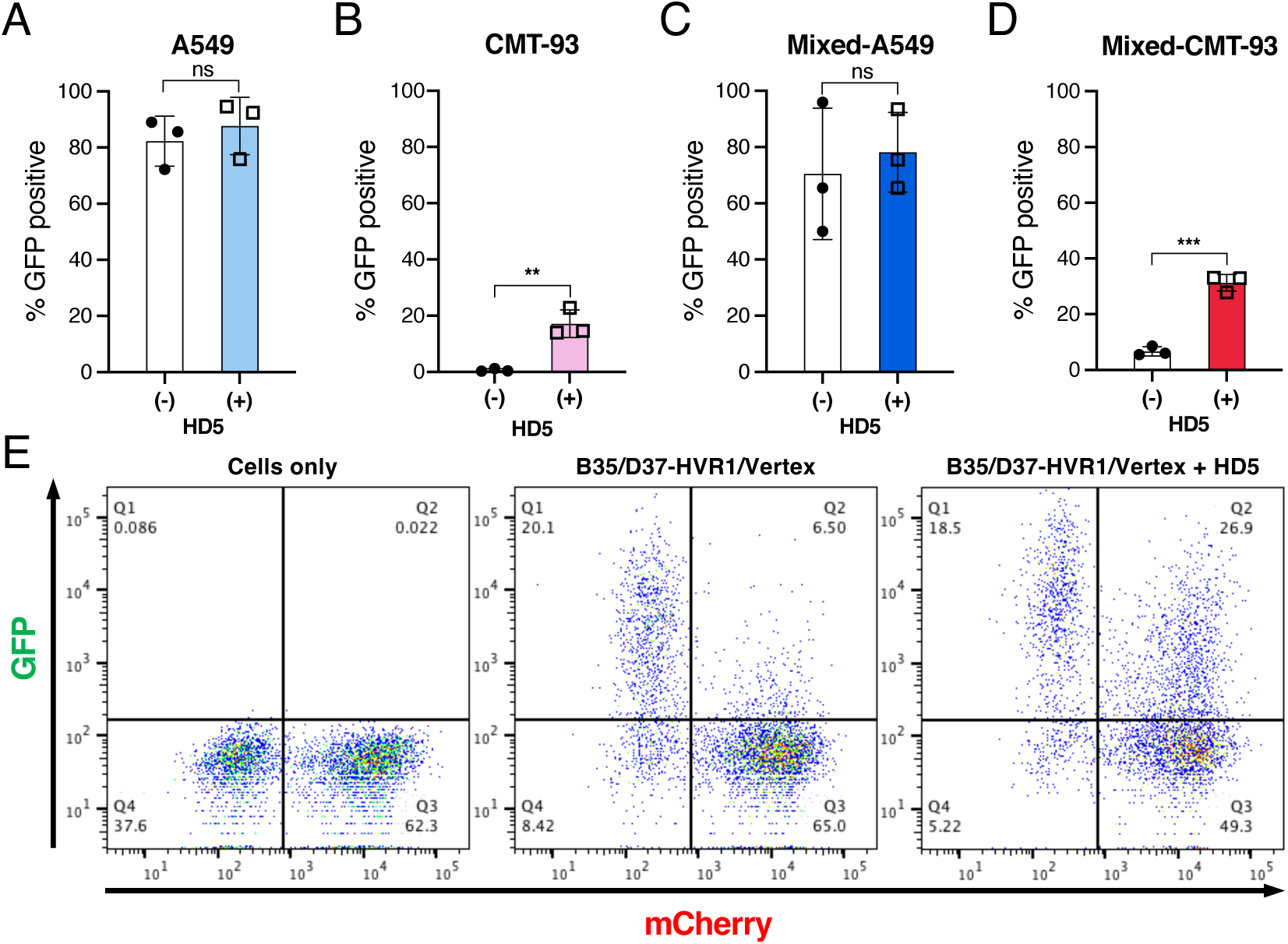
α-Defensin-mediated infection expands the tropism of HAdV. Infection of (A) A549 or (B) CMT-93 cells in monoculture or (C) A549 and (D) CMT-93 cells in mixed culture with B35/D37-HVR1/Vertex in the presence or absence of 20 μM HD5. Infected cells were enumerated by flow cytometry at 24 h post-infection. Data are the mean percentage of each cell type that are GFP positive ± SD of at least 3 individual experiments. (E) Representative flow cytometry plots of mixed cultures (left) without virus or infected with B35/D37-HVR1/Vertex in the (middle) absence or (right) presence of 20 μM HD5.

## Discussion

Our studies have revealed a novel pathway for HAdV binding to cells mediated by α-defensins. Unlike canonical HAdV receptor binding via the distal fiber knob, this mechanism bypasses the viral primary receptor. Accordingly, we found that it functions for multiple HAdVs that use different primary receptors (CAR, CD46, or sialic acid). However, our knowledge of the molecular interactions engaged by α-defensins to bridge the virus and cell is incomplete. Prior structural, biochemical, biophysical, and genetic evidence supports a direct interaction between α-defensins and all the major HAdV capsid proteins: fiber, penton base, and hexon [8, 10, 28, 31, 41, 46, 47]. These interactions contribute to defensin-mediated neutralization of susceptible serotypes. Yet, molecular determinants have only been defined for HD5 neutralization of HAdV-C5. Therefore, one important aspect of this study was to show that an HD5-enhanced HAdV-B virus could be engineered following principles derived from studies of HAdV-C. Nevertheless, the relationship between capsid features important for α-defensin neutralization of HAdV infection and those that mediate receptor-independent binding of HAdV to cells remains unresolved. It may be that distinct α-defensin binding sites on the capsid mediate either neutralization or cell binding but not both. More likely, there is some overlap, and perhaps one outcome (e.g., neutralization) requires more discrete determinants than the other. Regarding the other aspect of the bridging interaction, HD5 and HNP1 binding to cells, there are at least three possibilities: First, there may be one or more α-defensin-specific protein receptors. Receptors (e.g., CCR6) have been identified for the structurally related human β-defensins [48]. And inhibitor studies suggest that TLR4 signaling is important for increased transduction of human phagocytes by HAdVs in the presence of HNP1; however, direct binding to TLR4 either by HNP1 or by an HNP1/HAdV complex has not been shown [47]. In addition, the concentration of HNP1 (1 µM) that leads to increased phagocyte transduction has no effect in our assays. Thus, α-defensin receptors that might mediate the infection we have observed remain to be identified.

Second, most human α-defensins (HNP1, HNP2, HNP3, and HD5) function as lectins and bind to glycoproteins and glycolipids on the cell surface [39]. Third, α-defensins, being amphipathic, may interact directly with the phospholipid bilayer of the plasma membrane. Regardless of the type of molecular partner on the host cell, the interaction is preserved across species, since HD5 productively bridges interactions with mouse cells. Another critical aspect of the mechanism is the need for α-defensin multimerization, which we have shown for HD5 and may apply to HNP1 [28, 41]. We have previously demonstrated that the obligate monomer E21me HD5 can still bind to the HAdV capsid [28]. Therefore, we speculate that the requirement for HD5 self-association reflects an inability of a single defensin molecule to simultaneously bind both the virus and the cell. Thus, defensin-defensin interactions are an integral component of this bridging interaction. In summary, our studies have revealed a novel mode of HAdV binding to cells; however, important aspects of the virus-defensin, defensin-defensin, and defensin-cell interactions that underly this mechanism remain to be determined.

One critical insight from our studies is that although cell binding can be mediated by α-defensins, infection remains integrin-dependent. α-Defensins prevent the uncoating of neutralized serotypes, which become trapped in the endosome, and their genomes fail to reach the nucleus [5, 8, 28, 31, 49]. Rather than bypassing the clathrin-dependent entry pathway downstream of integrin-mediated internalization in which α-defensins exert their inhibitory activity, our data suggest that α-defensin-mediated entry of resistant viruses likely follows the same path. Therefore, our studies are consistent with a model in which the fate decision for infectivity is in the endosome and not on the cell surface.

For susceptible cells that express primary HAdV receptors, the α-defensin-mediated pathway likely contributes to enhanced infection that we have observed for certain HAdVs in the presence of HD5 and HNP1 [8, 10]. In these cells, we reason that both canonical receptor interactions and defensin-mediated binding function in parallel, converging on integrin co-receptor interactions. In support of this idea, we note that HAdV-D64 infection of A549 cells in the presence of HD5 approximates the sum of sialic acid-dependent and HD5-dependent infection. Moreover, C5/D64-HVR1 infection of A549 cells is enhanced by both HD5 and HNP1 while C5/D64-HVR1/RGE infection is not. However, we speculate that there could be additional effects of α-defensins that might increase cell binding and infection and contribute to enhancement. These include previously observed neutralization of the net-negative surface charge of the capsid that mitigates charge-charge repulsion at the cell surface and defensin-induced aggregation that alters the stoichiometry of virus-receptor interactions [28] as well as possible stabilization of viral protein interactions with canonical viral receptors. These principles may also apply to enhanced infection of mouse cells by MAdV-2 in the presence of mouse enteric α-defensins, which also bypasses competitive inhibition by fiber knob [9]. Assessing the relative importance of these pathways in different tissues and cell types will be an interesting direction for future study.

For non-susceptible cells, this new binding pathway leads to expanded tropism, allowing HAdV to enter cells either naturally lacking or engineered to be deficient in primary receptors. Moreover, HAdV infects non-susceptible cells through this pathway even in the presence of susceptible cells. α-Defensins are secreted into a milieu that often includes other cellular secretions (e.g., mucus) and microbiota, which could interfere with the bridging interactions required for this entry pathway. Nevertheless, α-defensin-mediated enhancement of both MAdV and HAdV occurs *in vivo*, suggesting that the necessary interactions can take place [9, 50]. Thus, even though we have not yet demonstrated expanded tropism in an animal model or a non-transformed cell culture model, we speculate that in the presence of sufficient concentrations of α-defensins, such as HNP1 in the lung and HD5 in the gut, integrin expression rather than primary receptor expression will dictate HAdV tropism *in vivo*.

Although receptor-independent binding of viruses to cells has never been previously described for α-defensins, there are other examples where host proteins bridge an interaction between HAdVs and cells. These proteins include lactoferrin and coagulation factors. For coagulation factors such as factor X, binding to hexon bridges interactions with heparan sulfate proteoglycans on the cell surface; however, this binding is not sufficient for internalization, which remains integrin-dependent for optimal uptake and infection [25]. Coagulation factor interactions are likely most important for injected adenoviral vectors for gene therapy or for natural infections that become viremic, which occurs rarely in immunocompetent individuals but can occur more commonly in immunocompromised individuals and may be fatal [51, 52]. In contrast, α-defensins and lactoferrin are found at mucosal surfaces, which are more commonly the sites of natural HAdV infection. Unlike α-defensin interactions, lactoferrin binding to the hexon of HAdV-C serotypes is sufficient to mediate both cell attachment and integrin-independent internalization into non-susceptible cells [26, 27]. However, as is true for α-defensins, the exact nature of the cellular receptor that mediates binding, how lactoferrin binding promotes efficient HAdV internalization, and the endosomal pathways that are then used are unknown. Collectively, interactions with these host molecules increase the ability of HAdVs to bind to cells and present more diverse strategies for viral evolution to drive tropism independently of fiber interactions. These interactions could also contribute to zoonotic infection, particularly since both α-defensin and lactoferrin-mediated entry of HAdVs have been demonstrated across species, although host range may then be restricted at other points in replication.

We speculate that the mechanism we have uncovered may apply to other viruses. In this regard, we have observed naturally α-defensin-resistant and –enhanced HAdVs, MAdVs, and rotaviruses, while AAVs can be resistant to but not enhanced by α-defensins [6–10]. If enhancement indeed represents the simultaneous functioning of receptor-dependent and defensin-dependent binding to cells, then the absence of enhancement may indicate that the defensin-dependent pathway is not productive for some viruses. The sufficiency of the defensin-dependent pathway likely depends on the nature of the cellular cues mediated by receptor interactions that trigger uptake and/or uncoating. And the essentiality of integrin co-receptors for HAdV infection indicates that binding to cells through defensin interactions alone does not signal for efficient uptake of these viruses. Therefore, although we have uncovered a new pathway for HAdV entry via hijacking of a host defensin molecule, whether it can be generalized as a mechanism of cell binding and entry for other viruses remains to be determined.

## Materials and Methods

### Viruses

The HAdV-C5-base vectors used in these studies are replication-defective, E1/E3-deleted, and contain a CMV promoter-driven enhanced green fluorescent protein (CMV-eGFP) reporter gene cassette in place of E1. The C5/D64-HVR1 was previously referred to as “Hexon HVR1 Chimera” in Diaz et al. [8]. The C5/D64-HVR1/RGE was generated by recombineering [53] the BACmid containing the C5/D64-HVR1genome to mutate the codons for RGD in the *penton base* gene to those for RGE. The replication-competent, E3-deleted, HAdV-D64-based vector containing a CMV-eGFP cassette in place of E3 was previously described [54]. The replication-defective, E1-deleted, HAdV-B35 vector with a substitution of *E4orf6* for that of HAdV-C5 and containing a CMV-eGFP cassette in place of E1 (HAdV-B35-ΔE1-CMV-GFP-E4orf6-HAdV-C5) was obtained from Silvio Hemmi (University of Zurich, Zurich, Switzerland) [55]. B35/D37-HVR1/Vertex was created by replacing the 3’ half of the *penton base* gene [base pair (bp) 14581–15378 in HAdV-B35, NCBI: AC_000019] with the 3’ half of HAdV-D37 *penton base* gene (bp 14371–15090, NCBI: AB448778), the EDES motif in HAdV-B35 fiber (encoded by bp 30878–30889) with the HAdV-D37 GYAR motif (encoded bp 31027–31038), and HVR1 of HAdV-B35 hexon (encoded by bp 18650–18745) with that of HAdV-D37 (encoded by bp 18199–18267) in sequential steps by recombineering the BACmid encoding HAdV-B35-ΔE1-CMV-GFP-E4orf6-HAdV-C5. The fidelity of all BACmid constructs was verified by Sanger sequencing of the recombineered region and by restriction digest of the entire BACmid. All adenoviruses were propagated in 293β5 cells, purified by CsCl gradient centrifugation, stored, and quantified as previously described [8].

### **α**-Defensin peptides

HNP1 and HD5 were created using peptides synthesized by CPC Scientific (Sunnyvale, CA) or LifeTein (Somerset, NJ), which were oxidatively re-folded, purified by reverse-phase high-pressure liquid chromatography (RP-HPLC) to homogeneity, lyophilized, resuspended in deionized water, and quantified by absorbance at 280 nm as described [7, 8]. Purity (>99%) and mass were verified by analytical RP-HPLC and mass spectrometry. Synthesis of E21me HD5 was described previously [40].

### Cells

A549 cells (ATCC) and CMT-93 mouse rectal carcinoma cells (a gift from Susan Compton, Yale University School of Medicine) were maintained in Dulbecco’s modified Eagle’s medium (DMEM) with 10% fetal bovine serum (FBS), penicillin, streptomycin, L-glutamine, and non-essential amino acids. Human haploid (HAP1) wild type and CD46 knockout (KO) cells were purchased from Horizon Discovery and cultured in Iscove’s Modified Dulbecco’s Medium (IMDM) with 10% fetal bovine serum (FBS), penicillin, and streptomycin. For each cell type, the appropriate basal media with additives and serum (complete medium) or as serum free media (SFM) was used in infectivity experiments.

Clonal CAR KO cells were created by targeting exon 2 of the *CXADR* gene in A549 cells for editing using a lentiviral vector encoding both the gRNA (5′-ACGCTTAGTCCCGAAGACCA-3′) and the *Streptococcus pyogenes* Cas9 nuclease. To create the lentiviral vector, 293β5 cells [56] were transfected with LentiCRISPR v2 (a gift from Feng Zhang, Addgene plasmid #52961, http://n2t.net/addgene:52961, RRID:Addgene_52961), pCMV-VSV-G (a gift from Bob Weinberg, Addgene plasmid #8454, http://n2t.net/addgene:8454, RRID:Addgene_8454), and psPAX2 (a gift from Didier Trono, Addgene plasmid #12260, http://n2t.net/addgene:12260, RRID:Addgene_12260) using Lipofectime 2000 (ThermoFisher Scientific). Vector-containing supernatant was harvested 72 h post-transfection and filtered through a 0.45 μm PES syringe filter. A549 cells transduced with this vector were selected with puromycin (1.5 µg/mL) and selection medium was changed every 3 to 4 days until all control cells died. Colonies were picked using cloning discs (Bel-Art) and transferred to individual wells of a 24-well plate containing medium without puromycin. Gene editing was assessed by sequencing a PCR product of the target region on an Illumina MiSeq. Sequence data was analyzed using CRISPResso2 [57].

CMT-93 cells expressing mCherry were created by transfection with pMCB320 (a gift from Michael Bassik, Addgene plasmid #89359, http://n2t.net/addgene:89359, RRID:Addgene_89359) using Lipofectamine 2000. Cells were selected in complete media containing with 3 µg/mL puromycin for 7 d prior to use in experiments.

All cells were tested at least quarterly for mycoplasma contamination.

### Surface expression of CAR

To confirm CAR ablation, A549 cells or CAR KO cells were trypsinized and fixed with 1% paraformaldehyde in PBS. After incubation with 20 mM glycine in PBS for 20 min, A549 cells (1.0 × 10^5^ cells/sample) suspended in PBS containing 0.2% sodium azide were stained with anti-CAR (clone RmcB, Millipore, USA) or mouse IgG1 isotype (Santa Cruz Biotechnology) antibodies at a concentration of 20 µg/mL followed by goat anti-mouse IgG (H+L) AF488 secondary antibodies (Invitrogen). Data was acquired on a BD FACS Canto II and analyzed using FlowJo v10.7.1.

### Quantification of viral infection

Monolayers of A549 cells were infected with serial dilutions of virus in black wall, clear bottom 96-well plates (PerkinElmer, San Jose, CA). Total monolayer fluorescence was quantified with a Sapphire (Azure) imager 24–30 h post-infection. For each viral stock, a concentration of virus producing 50–70% maximal signal was chosen for infectivity studies. The same concentration of virus was used on both A549 and CAR KO cells or both A549 and CMT-93 cells in parallel. For HAP1 and CD46 KO cells, a 10-fold dilution of viral stock was chosen.

To measure the effects of α-defensins on infectivity, purified virus was incubated with or without defensin for 45 min on ice in SFM. Confluent cells in black wall, clear bottom 96-well plates were washed two times with RT SFM, and virus/defensin mixtures were added in a final volume of 50 µL/well. After 2 h incubation at 37 °C, cells were washed two times with RT SFM and cultured with 100 µL complete media at 37 °C. Samples were incubated for approximately 24–30 h, washed with phosphate buffered saline (PBS), and scanned for eGFP signal using a Sapphire (Azure) imager. Fiji (version 2.1.0/1.53c) was used to quantify background-subtracted total monolayer fluorescence [58], and data are shown as a percent of control infection in the absence of defensin.

To block cell surface sialic acid, A549 cells were treated with 100 µg/mL wheat germ agglutinin (WGA) from *Triticum vulgaris* (Sigma) in 100 µL SFM for 1 h on ice at 4 °C and washed two times with SFM on ice prior to the addition of virus/defensin mixtures. To enzymatically remove sialic acid monosaccharides, A549 cells were treated with 10 mU/mL neuraminidase from *Arthrobacter ureafaciens* (Sigma) in 100 µL SFM for 1 h at 37 °C and washed two times with SFM on ice prior to the addition of virus/defensin mixtures.

For mixed culture infections, the same number of mCherry expressing CMT-93 cells alone, A549 cells alone, or an equal mixture of A549 and mCherry expressing CMT-93 cells were seeded at 60,000 cells per well in 48-well plates and incubated overnight at 37 °C. Infections were carried out as described above except that virus/HD5 mixtures were added in 100 µL/well, and 250 µL/well complete media was added after washing. After incubation for 24– 48 h, cells were trypsinized, fixed with 1% paraformaldehyde in PBS, and analyzed on a BD FACS Canto RUO. The percentage of GFP positive cells was calculated independently for the mCherry negative (A549) and mCherry positive (CMT-93) cells in each sample. Note that CMT-93 cells were 91-98% mCherry positive, and any mCherry-negative CMT-93 cells were indistinguishable from A549 cells in this assay.

### Quantification of virus bound to cells

Viruses were labeled with Alexa Fluor 488 (AF488) carboxylic acid, tetrafluorophenyl ester (Thermo Fisher Scientific) as previously described [31]. Labeling had less than a two-fold effect on viral infectivity on A549 cells. Viruses were incubated with or without HD5 for 75 min on ice in 60 μL SFM. A549 cells (1.0 × 10^5^ cells/sample) were trypsinized and incubated in suspension on ice in PBS containing 0.2% sodium azide to prevent endocytosis. The virus/defensin mixtures were added to the cells (final volume, 100 μl/sample), incubated on ice for 45 min, washed two times with cold PBS, and fixed with 1% paraformaldehyde in PBS. Samples were analyzed on a BD FACS Canto II, and a geometric mean fluorescence (GMF) value for the AF488 signal of the entire cell population for each sample was calculated using FlowJo v10.7.1. Note that although a similar concentration of each virus was used, the absolute shift in GMF varied because each virus was labeled to a different level of brightness. Fold change in GMF was calculated relative to virus bound to WT cells in the absence of HD5 or E21me HD5 for each virus.

### Statistical analysis

Statistical tests were performed using Prism 10.0.2 as follows: Figures 1B and 1C: Repeated measures one-way ANOVA with the Geisser-Greenhouse correction comparing each condition to control infection of untreated cells in the absence of HD5, with Dunnett’s multiple comparisons test and individual variances computed for each comparison. Figures 2E and 4A: Repeated measures two-way ANOVA comparing each condition to the amount of virus bound to WT cells in the absence of HD5, with Dunnett’s multiple comparisons test with a single pooled variance. Because the absolute fluorescent labeling of all viruses was not uniform, samples were compared as a separate family for each virus, and the GMF of cells alone was different relative to control cells for each virus. For Figures 3B, 4C, 4D, and 5B: Repeated measures two-way ANOVA comparing C5/D64-HVR1 to C5/D64-HVR1/RGE at each defensin concentration, with Šídák’s multiple comparisons test with a single pooled variance. For CAR KO cells, log-transformed values were compared. Note that WT HAdV-C5 data is included on these graphs for illustrative purposes but was not compared to either chimeric virus. For Figure 6C: Paired, two-tailed t test of the log-transformed data. For Figures 7A to 7D: Unpaired, two-tailed t test. For all graphs, significance is marked by asterisks: *, P = 0.01 to 0.05; **, P = 0.001 to 0.01; ***, P = 0.0001 to 0.001; ****, P < 0.0001. Unmarked comparisons within the specified analysis range are not significant (P > 0.05). Note that for data analyzed and graphed on log scales, a value of 0.1 µM was imputed for the absence of HD5, and a value of 0.0156% was imputed for no infection.

## Acknowledgements

We thank Danielle R. Williams for assistance with the creation of CAR KO A549 cells. This work was supported by R01 AI104920, from the National Institute of Allergy and Infectious Diseases (www.niaid.nih.gov) and by the Office of the Director, National Institutes of Health (www.nih.gov/institutes-nih/nih-office-director) under Award Number S10 OD026741 (to J.G.S.). Additional support to J.G.S. in the form of subsidized core services was provided by NIH grants UL1 TR000423 from the National Center for Advancing Translational Sciences (ncats.nih.gov), and P30 CA015704 from the National Cancer Institute (www.cancer.gov). The funders had no role in study design, data collection and analysis, decision to publish, or preparation of the manuscript.

